# The study on the mechanism of “oil-spilling” of wolfberry (Lycium barbarum) fruits during storage

**DOI:** 10.1101/492272

**Authors:** Fu Wang, Suqin Xiong, Youping Liu, Lin Chen, Hongping Chen

**Author notes:** Corresponding author: Hongping Chen and Lin Chen.

## Abstract

Improper storage of wolfberry fruit is liable to soften, discoloration and adhesion, we call it “oil-spilling”. The purpose of the study is to explore the mechanism of “oil-spilling” of wolfberry fruits during storage. In our study, the relationship between water content, superoxide dismutase (SOD), peroxidase (POD) activity, malondialdehyde (MDA) content and the cross section, epidermal cells of wolfberry fruit were studied during storage. Results showed that wolfberry fruits began to “oil-spilling” on the 20th day, and with the extension of storage time, the degree of “oil-spilling” gradually deepened, the content of water increased and the activity of POD decreased, and the content of MDA increased. It suggested that “oil-spilling” was the result of improper external storage conditions, which it could promote the increase of water content and reactive oxygen free radical of wolfberry fruit, and lead to damage of cell structure and function and lipid peroxidation of cell membrane. The results explain the mechanism of “oil-spilling” and provide a strong basis for the establishment of scientific storage methods of wolfberry fruit.

## 1 Intoduction

*Lycium barbarum* berries, also named wolfberry fruits, have been used in China and other Asian countries for more than 2000 years as a traditional medicinal herb and food supplement. It is also widely used in food and health care industry all over the world because of its obvious beneficial effects including neuroprotection, anti-aging and anticancer properties, and immune modulati^[1,2]^. However,wolfberry fruit is often accompanied by easy to soften, mutual adhesion, and color-changing during storage, we call it “oil-spilling”^[3]^. The occurrence of this phenomenon will seriously affect the character, chemistry, utility quality and clinical drug safety of wolfberry fruit^[4]^. At present, the scientific research on storage and maintenance of Wolfberry fruit was mainly focused on the effects of external storage conditions, packaging materials on the quality of medicinal materials and changes in the contents of main active components before and after deterioration^[5,6,7,8]^. However, the mechanism of “oil-spilling” has not been reported.

Our research group found that the change of water content was closely related to the time and degree of “oil-spilling”^[10]^. So this paper put forward the hypothesis of the mechanism of “oil-spilling”of wolfberry fruit based on the plant active oxygen free radical senescence hypothesis^[9]^. The water in wolfberry fruit accelerates the cell metabolism rate, thus produces excessive reactive oxygen free radical to destroy the cell structure and function, and then leads to the increase of cell membrane permeability, than the phenomenon of “oil-spilling” occurs. To verify this assumption, this study placed wolfberry fruit in the drug stability test box referred to the “Traditional Chinese medicine, natural drug stability research technical guidelines”. Methods of sampling at different time points were used to study the relationship between water content, superoxide dismutase(SOD), peroxidase(POD) activity, malondialdehyde (MDA) content and the cross section and epidermal cells of wolfberry fruit during storage. The objective of this study is to explain the mechanism of “oil-spilling” of wolfberry fruit during storage and to provide a theoretical basis for the scientific storage of wolfberry fruit.

## 2 Material and methods

### 2.1 Materials

The fruit of wolfberry fruit was purchased from Zhongning County, Jingyuan County, Yinchuan City, Ningxia Province. It was identified as Lycium barabarum L. by Professor Lu Xianming, Chinese Medicine specimen Center of Chengdu University of traditional Chinese Medicine.

### 2.2 Instruments and reagents

Nanodrop2000 Spectrometer (Thermo Fisher Scientific); BP211D Electronic Analytical balance (Sartorius, Germany); Ultraviolet Spectrophotometer Agilent (8453) (Agilent, USA);SB25-12D Ultrasonic Cleaner (Ningbo Xinyi Ultrasonic equipment Co., Ltd.); WHP150 drug stability test box (Chongqing InBev Experimental instrument Co., Ltd); Haier Vertical refrigerator (SC-316); TGL-16 desktop high-speed refrigeration centrifuge ( Hunan Xiangyi Laboratory instrument Development Co., Ltd.); Tissuelyser-48 automatic sample lapping instrument (Shanghai Jingxin Test equipment Science and Technology Ministry); 48T malondialdehyde (MDA) test kit (Nanjing Institute of Biological Engineering, No.: A001-2); SOD kit (side typing) (Nanjing Institute of Bioengineering, No.: A084-3);POD kit (Plant) (Nanjing Institute of Bioengineering, No.: A003-3).

### 2.3 Methods of storage and sampling

Three kinds of wolfberry fruit were stored in constant temperature and humidity box (temperature: 40 °C, humidity: 75%). Sampling and detection were carried out on the day of 0d, 20d, 40d and60d, respectively, and the results were shown in Fig 1.

**Fig 1.**
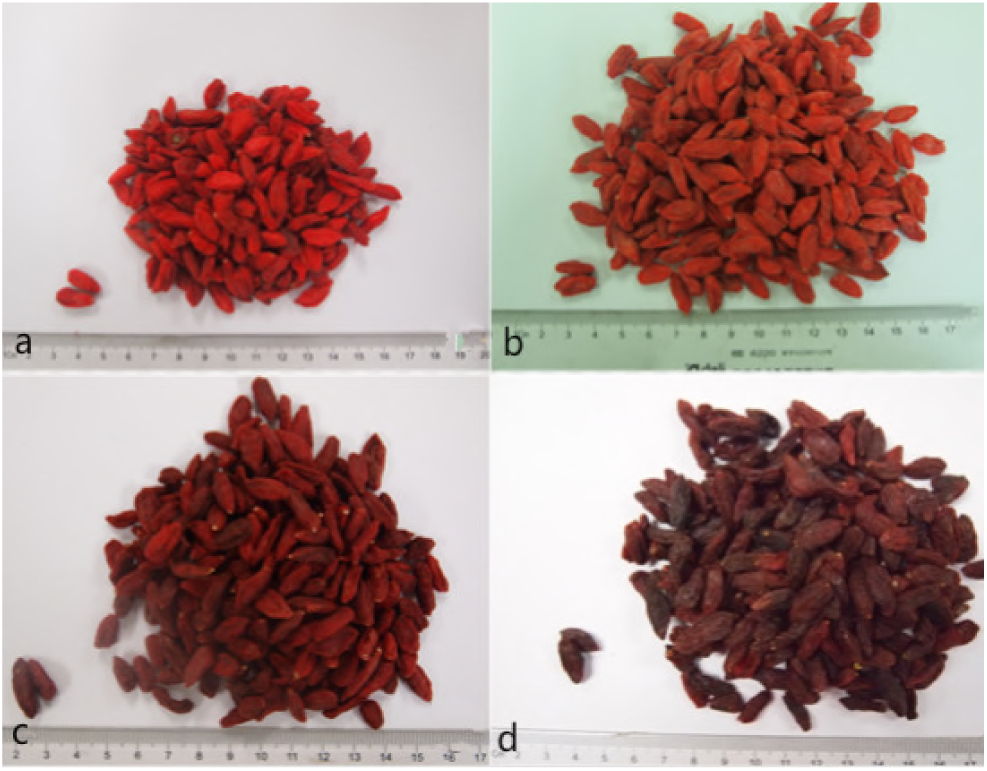
Sampling pictures of wolfberry fruit at 0day(a),20days(b),40days(c)and60days(d)

### 2.4 Determination of polysaccharides content at different Storage time

Through the preliminary study of this subject, Polysaccharide is the main active component of wolfberry fruit, and the content of polysaccharide change obviously in the process of “oil-spilling. Therefore, the content of polysaccharides was selected as the main index component in this experiment. Methods of determination refer to Yang et al^[11]^.

### 2.5 Determination of water content under different storage time

The results of the previous study showed that the water content of wolfberry fruit was closely related to the physiological function of plant tissue. Therefore, the change of water content in wolfberry fruit during storage was studied in this study. Methods of water content determination refer to Sarkar et al^[12]^.

### 2.6 Determination of superoxide dismutase (SOD) under different storage time

Accurately weighed the wolfberry sample 0.1g, put the sample into liquid nitrogen and fully grind, added 9 times of normal saline and mix with a vortex meter. The sample was then centrifuged in a cryopreservation centrifuge, and the supernatant 0.1ml was added to the peroxidase kit reaction reagent. The sample was accurately reacted for 30 minutes in a water bath at 37 °C. Finally, the reaction termination reagent was added and the supernatant was determined by centrifugation at 420nm^[13]^.

### 2.7 Determination of peroxidase (POD) content under different storage time

Accurately weighed the wolfberry sample 0.1g, put it into liquid nitrogen and fully grind, added 9 times of normal saline and mix with a vortex meter, then the sample was centrifuged in a cryopreservation centrifuge, 30 μl supernatant was taken, superoxide dismutase reagent was added into the reagent, and the reaction time was 40 min at 37 °C water bath. Finally, the chromogenic agent was added and statically for 10 min. The results were determined at 550nm^[14]^.

### 2.8 Determination of malondialdehyde (MDA) content under different storage time

Accurately weighed the wolfberry sample 0.1g, put it into liquid nitrogen and fully grind, added 9 times of normal saline and mix with a vortex meter. The sample was then centrifuged in a cryogenic centrifuge, 50 μl supernatant was added to the working fluid of malondialdehyde (MDA) kit and reacted for 20 min in a metal bath above 95 °C. Then the sample was cooled with flowing water and 0.25ml was added to the 96-well plate. The enzyme-labeled solution was used for determination at 530nm^[15]^.

### 2.9 Observation on transverse section of wolfberry fruit tissue

PAS stain:Fresh tissue samples were taken and put into stationary solution (70% alcohol: glacial acetic acid: formaldehyde 90: 5: 5), then dehydrated in different concentrations of alcohol,impregnated with wax and embedded. The conventional slice was 10-12 μm, treated with dewaxing, and then added 0.15% periodate to oxidize 5~10min. Rinsed with evaporated water for 2 times and Dyeing 10min with Svhiff solution without Light; 15% heavy sulfurous acid steel was washed twice, each 2~5min;Harris hematoxylin was soaked for 2 min and washed with tap water for 5 min; Soaked in 1% hydrochloric acid alcohol, then rinse with double steamed water. Finally, it was fixed with xylene transparent neutral gum^[18]^.

HE stain:Fresh tissue samples were taken and put into stationary solution (70% alcohol: glacial acetic acid: formaldehyde 90: 5: 5), then dehydrated in different concentrations of alcohol. Then the tissue block was placed in xylene for transparent treatment, so that xylene can gradually replaced the ethanol solvent in the tissue. Placed the transparent tissue blocks in the hexagonal paraffin wax for wax immersion, and then put them in the wax dissolution box for heat preservation. After the paraffin was fully immersed in the tissue block, pour the hexagonal paraffin into the tissue and quickly clip the paraffin-soaked tissue. After cooling and solidifying into blocks, the sections can be stained^[17]^.

The epidermis cells were also observed by microscope^[16]^.

## 3 Results and discussion

### 3.1 Changes of water content in wolfberry fruit

As shown in Table 1, the water content of wolfberry fruit increased with the storage time. And the degree of “ oil-spilling” has been deepened with the increase of water content, which indicated that there was an obvious correlation between the water content and “ oil-spilling”. Water content is of great significance to the safe storage of wolfberry fruit. Liu Jiayu et al ^[24]^studied the correlation between appearance character and water content of wolfberry fruit. The results showed the phenomenon of color-changing appeared when the water content of wolfberry fruit was more than 13%, and the phenomenon of “oil-spilling” appeared when the water content was more than 18%. Therefore, the water content of wolfberry is one of the key factor of “oil-spilling”.

**Table 1.**
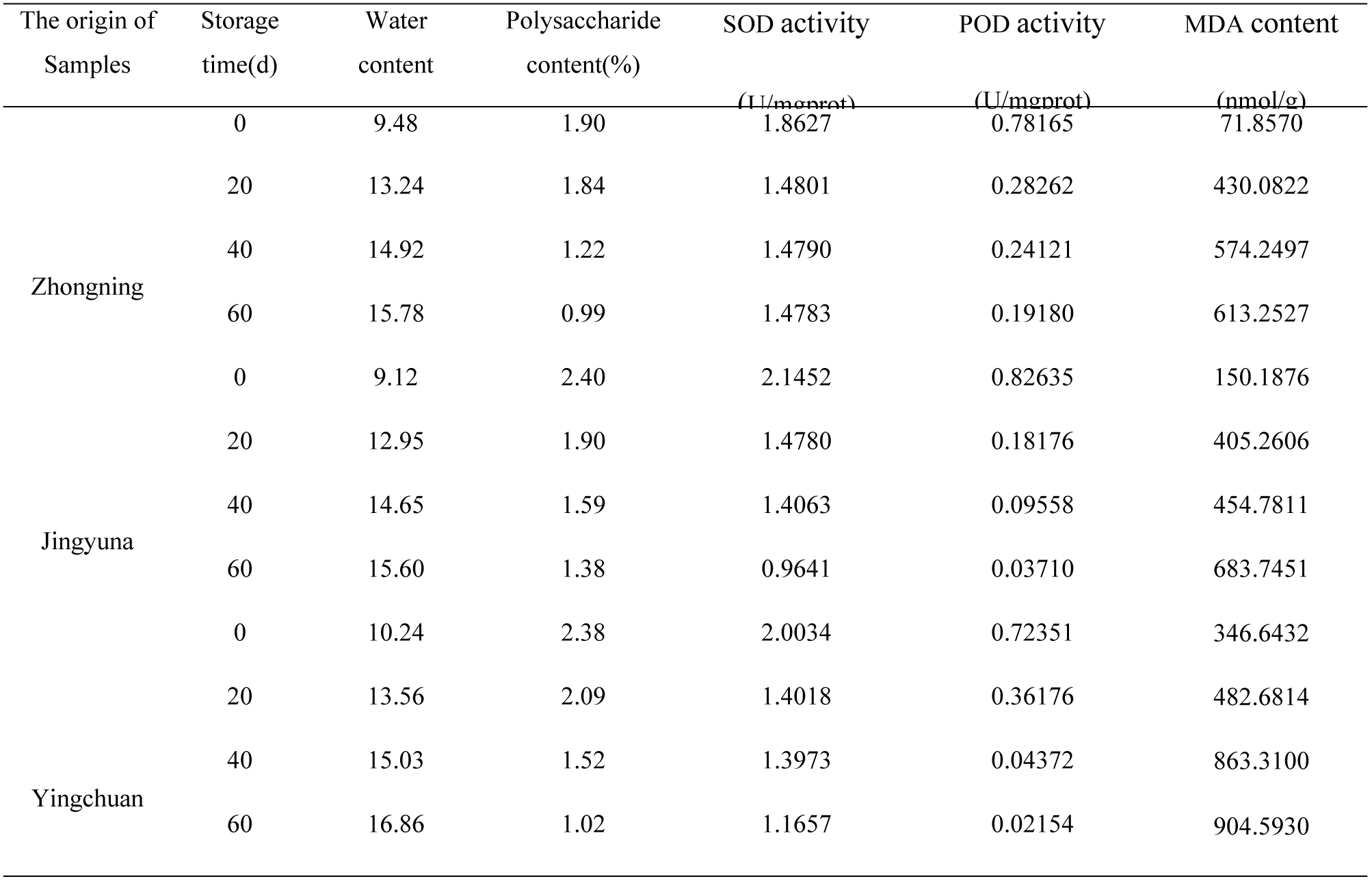
Water, MDA content and SOD, POD activity of wolfberry fruit

### 3.2 Changes of MDA content and SOD, POD activity in wolfberry fruit

Table 1 shows the changes of MDA content and SOD, POD activity in wolfberry fruit. The content of MDA increased significantly with the prolongation of storage time in all the three batches of samples, SOD and POD activity of wolfberry fruit decreased.

SOD and POD can oxidize the harmful substances in the cells, so it has protective effect on the cells^[23]^. MDA is one of the products of peroxidation, which can react with various components of the cell, thus causing serious damage to enzymes and membranes, reducing membrane resistance and membrane fluidity, and ultimately destroying the structure and physiological integrity of the membrane^[22]^. Dhindsa et al^[21]^ studied the changes of SOD activity during senescence of oats in vitro and found that the activity of SOD decreased and the content of MDA increased. In this study, the “oil-spilling” of wolfberry fruit was studied during different storage time. The results shown that the content of MDA increased and the activity of POD and SOD decreased with the increase of the degree of “oil-spilling” of wolfberry fruit. There is an enzyme protection system for scavenging oxygen free radicals in wolfberry fruit cells^[20]^. SOD and POD can keep the active oxygen free radicals at a certain level, thus avoiding or reducing the damage of free radicals to biological macromolecules^[19]^. Therefore, when the activity of POD or SOD decreases and the content of MAD increases in wolfberry fruit, it will cause serious damage to the cell membrane, and the contents of polysaccharides in the cell will overflow gradually, which will eventually lead to the phenomenon of “oil-spilling”.

### 3.3 Changes of polysaccharides content in wolfberry fruit

Results of the content of polysaccharides are shown in Tabel 1 and Fig 5. The content of polysaccharides in all the three samples decreased significantly, and with the prolongation of storage time, the content of polysaccharides decreased gradually and the degree of “oil-spilling” deepened. The changes of MDA content and SOD, POD activity in wolfberry fruit has been verified the structure and function of the cell were destroyed, and the membrane lipid peroxidation and permeability was increased. As a result, polysaccharides were exosmated and the content decreased. Polysaccharide is the main active component of wolfberry fruit^[25,26]^, and the quality will be seriously affected by the decrease of polysaccharide content. Therefore, the occurrence of “oil-spilling” has a negative impact on its quality.

### 3.4 Changes of Cross section and epidermal cells in wolfberry fruit

From figure 2, as the degree of “oil-spilling”of wolfberry fruit became more and more serious, the color of cell wall and cell membrane of the cross section pulp dyed by HE gradually desalinated. At the same time, the number of stained starch granules was gradually reduced in the flesh cells of the cross section after PAS reagent dyeing (Fig 3), but the golden yellow oil particles in the epidermal cells were increasing (Fig 4). The results showed that with the deepening of the “oil-spilling”of wolfberry fruit, the degree of cell structure damage was also increasing. As a result, the contents of pulp cells were constantly infiltrated into epidermal cells, and the golden yellow oily granules in the cells increased continuously. The results were in agreement with the exosmosis phenomenon of “oil-spilling”.

**Fig 2.**
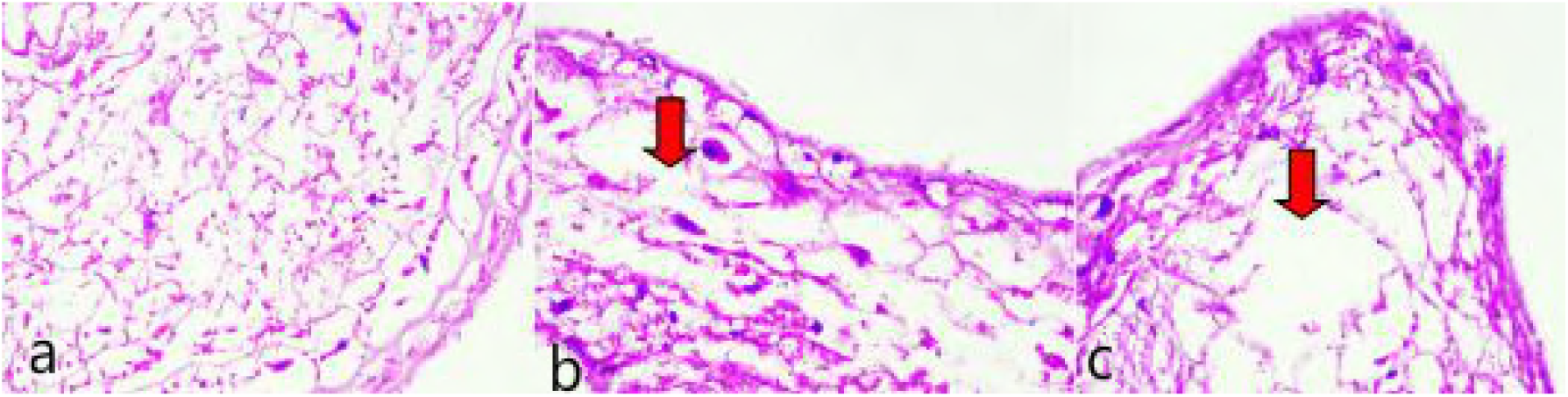
Cross sectional HE staining of wolfberry fruit at 14 days(a),22 days(b)and 30 days(c)

**Fig 3.**
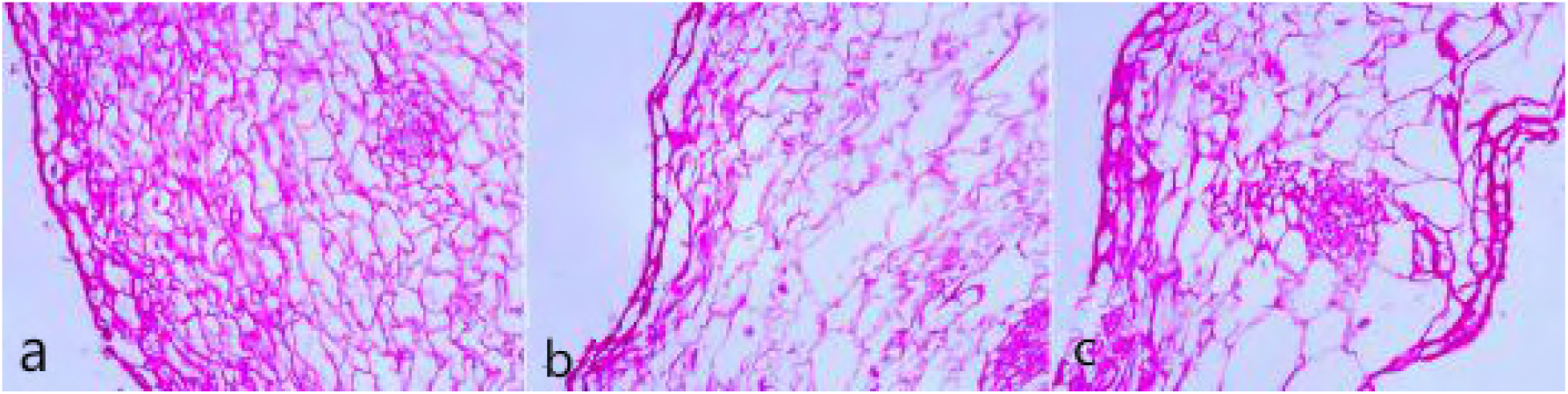
Cross sectional PAS staining of wolfberry fruit at 14 days(a),22 days(b)and 30 days(c)

**Fig 4.**
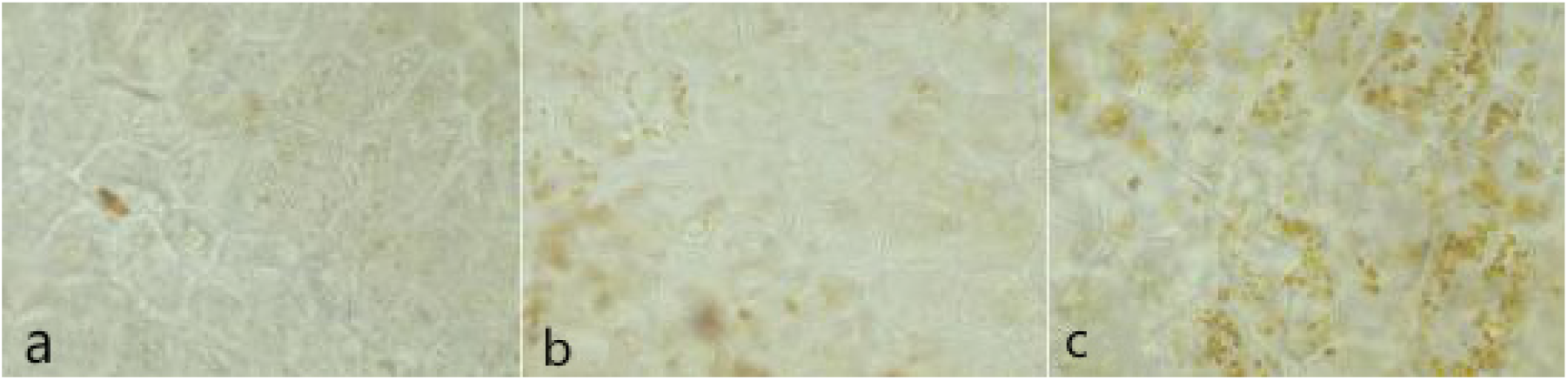
The changes of epidermis cells of wolfberry fruit at earlier stage(a),mid-term stage(b)and later stage(c) of “oil-spilling”

**Fig 5.**
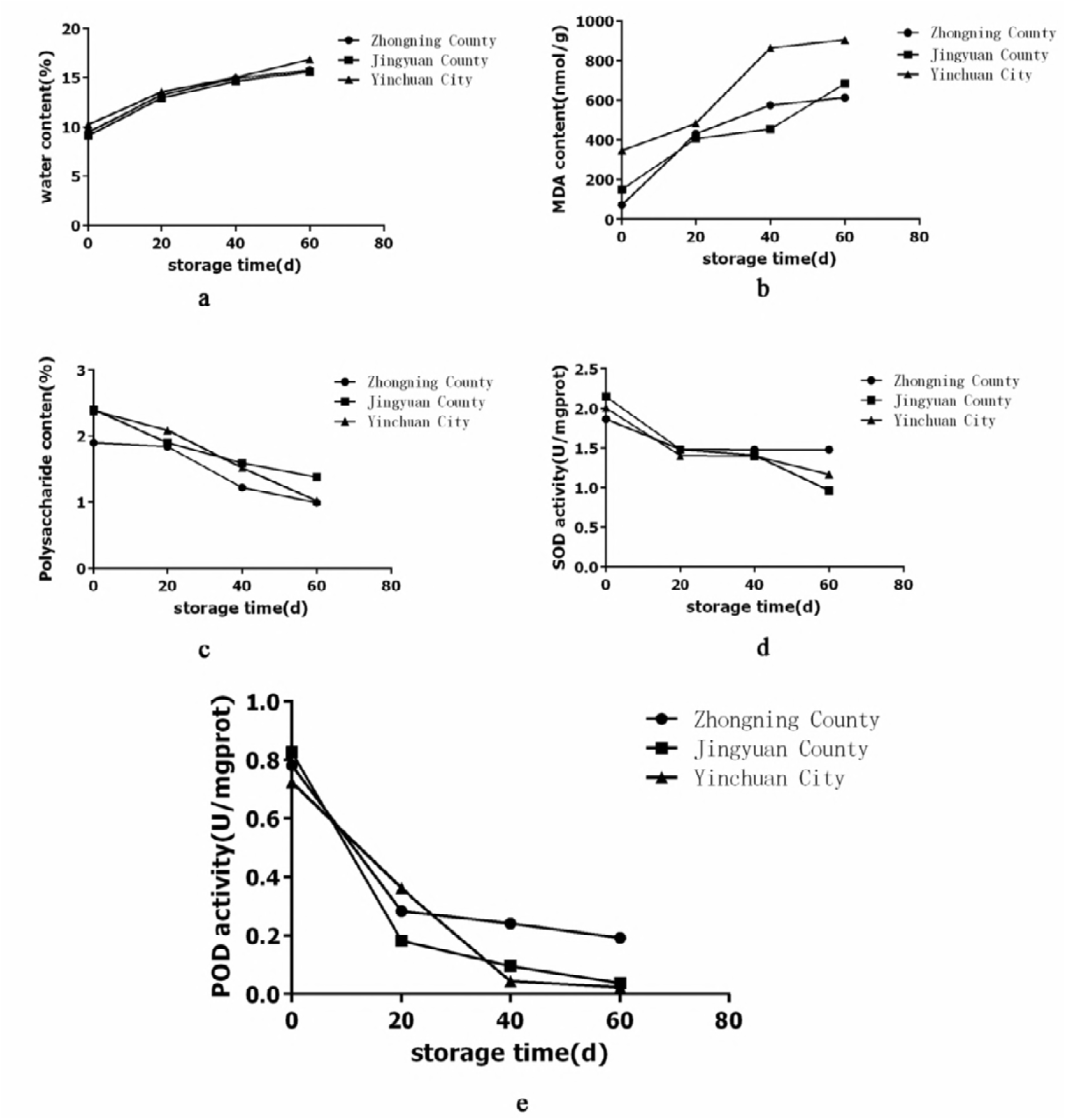
The content of water(a), MDA(b), Polysaccharide(c) and the activity of SOD(d), POD(e)

**Fig 6.**
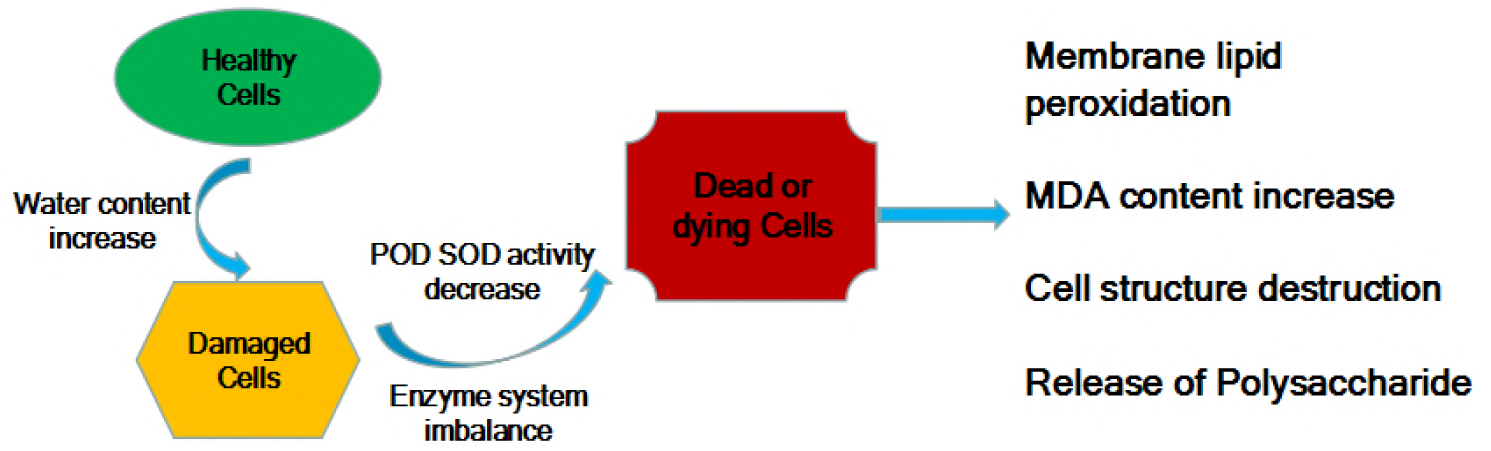
The mechanism of “oil-spilling” of the cells of wolfberry fruit

## 4 Conclusion

In this paper, the hypothesis of the mechanism of “oil-spilling” was put forward from the point of view of the change of intrinsic physiological activity of wolfberry fruit, and the content of polysaccharide and the water content were measured in different storage periods. Activity of reactive oxygen free radical scavenging enzyme and the content of malondialdehyde were also determined. The results showed that the activity of active oxygen free radical scavenging enzyme SOD, POD in wolfberry fruit decreased with the prolongation of storage time, and the cell structure and function were destroyed, which resulted in membrane lipid peroxidation. As a result, malondialdehyde (MDA) content continues to increase, leading to the occurrence of “oil-spilling”.

The results of polysaccharide and water determination showed that the wolfberry fruit appeared “oil-spilling” on the 20th day, and the content of polysaccharides decreased obviously, which indicated that the occurrence of “ oil-spilling” was the result of improper storage. It is suggested that the storage conditions should be paid attention. At the same time, high temperature and high humidity should be avoided.

## Acknowledgements

This research was supported by the National Natural Science Foundation of China(8140140928). The authors thank Zhuyun Yan for technical assistance.

## Author Contributions

Conceived and designed the experiments:HCP LC. Performed the experiments:FW SQX. Analyzed the date:YPL. Wrote the paper:FW.

